# The gut microbiota affects the social network of honeybees

**DOI:** 10.1101/2021.12.31.474534

**Authors:** Joanito Liberti, Tomas Kay, Andrew Quinn, Lucie Kesner, Erik T. Frank, Amélie Cabirol, Thomas O. Richardson, Philipp Engel, Laurent Keller

## Abstract

The gut microbiota influences animal neurophysiology and behavior but has not previously been documented to affect emergent group-level behaviors. Here we combine gut microbiota manipulation with automated behavioral tracking of honeybee sub-colonies to show that the microbiota increases the rate and specialization of social interactions. Microbiota colonization was associated with higher abundances of one third of metabolites detected in the brain, including several amino acids, and a subset of these metabolites were significant predictors of social interactions. Colonization also affected brain transcriptional processes related to amino acid metabolism and epigenetic modification in a brain region involved in sensory perception. These results demonstrate that the gut microbiota modulates the emergent colony social network of honeybees, likely via changes in chromatin accessibility and amino acid biosynthesis.

## Main

Understanding which factors regulate the organization of animal societies is a long-standing goal of biological research (*1*). While various genetic and ecological factors have been associated with the diversity of social organization across the animal kingdom (*2-5*), much less is known about the role of symbiotic interactions with the gut microbiota, which is increasingly recognized as an important modulator of neurophysiology. Bacterial metabolites and signals produced in the gut can reach the brain and elicit local host responses that affect the host nervous system (*6-8*). There is also accumulating evidence linking the gut microbiome to social behavior and its dysfunctions (*9-12*). However, effects on social behavior have typically been investigated during one-to-one encounters between gnotobiotic animals, and in model organisms that do not naturally express complex social structure (*7, 8, 13-17*). Whether and how the diversity of gut microbes hosted by individual animals influences the emergent properties of group living remains unknown.

To address this question, we performed controlled laboratory experiments with honeybees, which live in complex societies and exhibit division of labor among colony members (*18, 19*). In these societies, individuals follow simple decision-making strategies to produce elaborate social phenotypes at the colony-level. Honeybees provide a powerful model to explore how the gut microbiota affects group-level social phenotypes for several reasons. First, they exhibit complex but experimentally tractable social behavior (*20, 21*). Second, they have a well-characterized and evolutionarily stable gut microbiota comprising relatively few species (*22-24*). Third, the composition of this community can be manipulated and the resulting gnotobiotic bees (i.e., with defined microbiota) studied under controlled lab conditions (*16, 23, 24*). Finally, the gut microbiota has been shown to affect hormonal signaling, gene expression of insulin-like peptides in the head, and sugar intake (*25*), indicating substantial crosstalk between the gut and the brain along what is referred to as the gut-brain axis.

To investigate the influence of the gut microbiota on bee behavior and colony social organization we produced gnotobiotic bees to be either microbiota-depleted (MD) or colonized with a homogenate of five nurse bee guts (CL) reconstituting the natural gut microbiota (*25-27*). We used an automated behavioral tracking system (*28, 29*) to monitor social interactions for a week in nine pairs of sub-colonies of ∼100 three-day-old worker bees (Fig. 1A and B and Supplementary Movie 1). The microbiota treatment led to clear differences in the abundance and taxonomic composition of the gut bacterial communities (Figs. S1 and S2; Permutational multivariate analysis of variance (PERMANOVA) using Bray-Curtis dissimilarities calculated from a matrix of absolute abundances of amplicon sequence variants (ASVs): *F*_1,179_=65.99, *R*^2^=0.27, *P<*0.001). Bees in both treatments had a circadian rhythm and a pattern of interactions reflecting natural behavior (Figs. 1C and S3). The microbiota treatment had a significant effect on behavior, with CL bees having a higher rate of head to head interactions than MD bees (Fig. 1C and D; paired *t*-test: *t*=2.82, df=8, *P*=0.02). CL bees also exhibited a much higher degree of specialization (measured by the standard deviation in edge-weight per node in the social network) than MD bees (Fig. 1E; paired *t*-test: *t*=2.93, df=8, *P*=0.02) suggesting that CL bees formed stronger social ties with specific subsets of nestmates, while MD bees interacted more randomly within the colony. Importantly, these differences did not simply reflect a treatment effect on overall activity level. First, there was no significant difference between CL and MD bees in the rate of contacts (interactions not involving the head of bees) (Fig. 1F; paired *t*-test: *t*=0.32, df=8, *P*=0.76). Second, CL and MD bees exhibited similar movement patterns (average speed and within-individual variation in speed; Figs. 1G and S4A). And third, the gut microbiota had no significant effect on survival (Fig. S4B; paired *t*-test: *t*=1.10, df = 8, *P*=0.30). Hence, these results suggest that the gut microbiota specifically promotes and structures social interactions.

**Figure 1:**
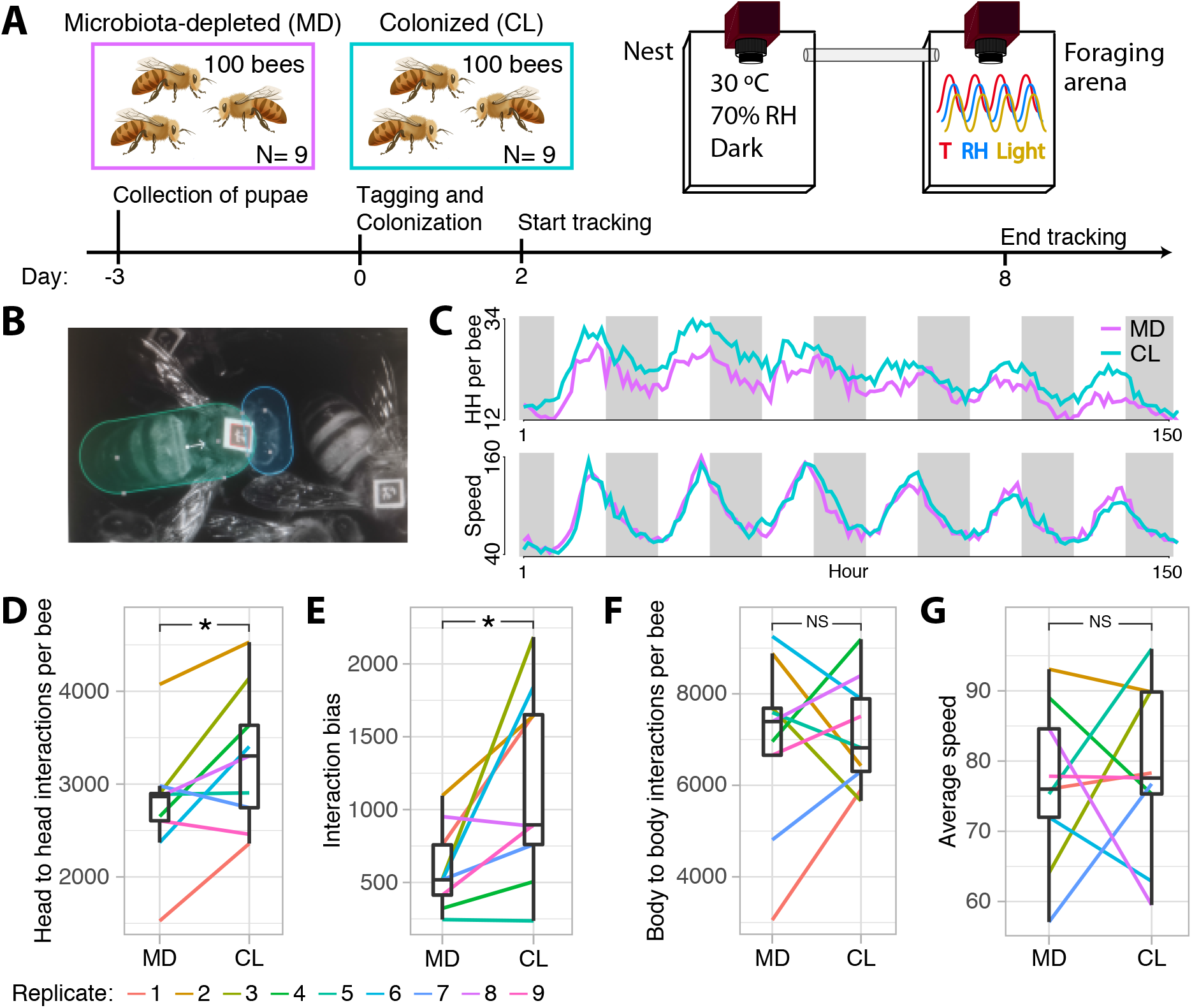
The gut microbiota affects honeybee social behavior. (A) Experimental design and timeline for a single experimental replicate. Gnotobiotic bees were produced by rearing pupae in an incubator and colonizing them with their treatment solution as newly emerged adults. Each sub-colony of ∼100 bees could move freely between two plexiglas boxes hosted within separate climate-controlled chambers. Social interactions were quantified by monitoring the orientation and position of individual tags glued onto the thorax of each bee and (B) counting overlaps between ellipses drawn over the bees’ heads and bodies. (C) Line plots showing the number of head to head interactions (HH per bee) and average speed per hour during 152 h of tracking averaged across all experimental replicates, and colored by gut microbiota treatment. White and grey shading represent day and night, respectively. (D) Average number of head to head interactions per bee for each sub-colony during the week of tracking. (E) Interaction bias, representing average variance in head to head interactions per bee per sub-colony. (F) Average number of body to body contacts per bee per sub-colony. (G) Average speed per bee per sub-colony. Lines connect paired colonies in each experimental replicate. Box plots show the median and first and third quartiles. Whiskers show the extremal values within 1.5 times the interquartile ranges above the 75th and below the 25th percentile. * *P<*0.05, NS = not significant.

To probe how the microbiome may affect social behavior, we analyzed soluble metabolites in the brain and hemolymph of a random subset of bees across the experimental replicates (brain, n=167; hemolymph, n=159). More than a third (21/60) of the metabolites detected in the brain differed significantly in abundance between MD and CL bees (BH-adjusted *P*<0.05) (Fig. 2A and Table S1). Strikingly, all of the differently abundant metabolites were more abundant in CL than in MD bee brains, and there was an over-representation of amino acids and intermediates of amino acid metabolism (Fig. 2C and Table S2; Fisher exact test, *P*= 0.031). CL bees had a higher abundance of three out of the six essential and eight out of the 15 non-essential amino acids (*30*), as well as three out of the seven metabolites linked to amino acid metabolism (Fig. 2C and Table S2). Several of the differently abundant amino acids (e.g. serine, glutamine, aspartate, glycine) have known roles in synaptic transmission and brain energetic function (*31, 32*). This pattern in the brain contrasted with that of the hemolymph, where less than 8% (6/76) of the metabolites were significantly differently abundant between MD and CL bees, including three that were also differently abundant in the brain (glutamine, 5-oxoproline, and an unidentified metabolite; Fig. 2B).

**Figure 2:**
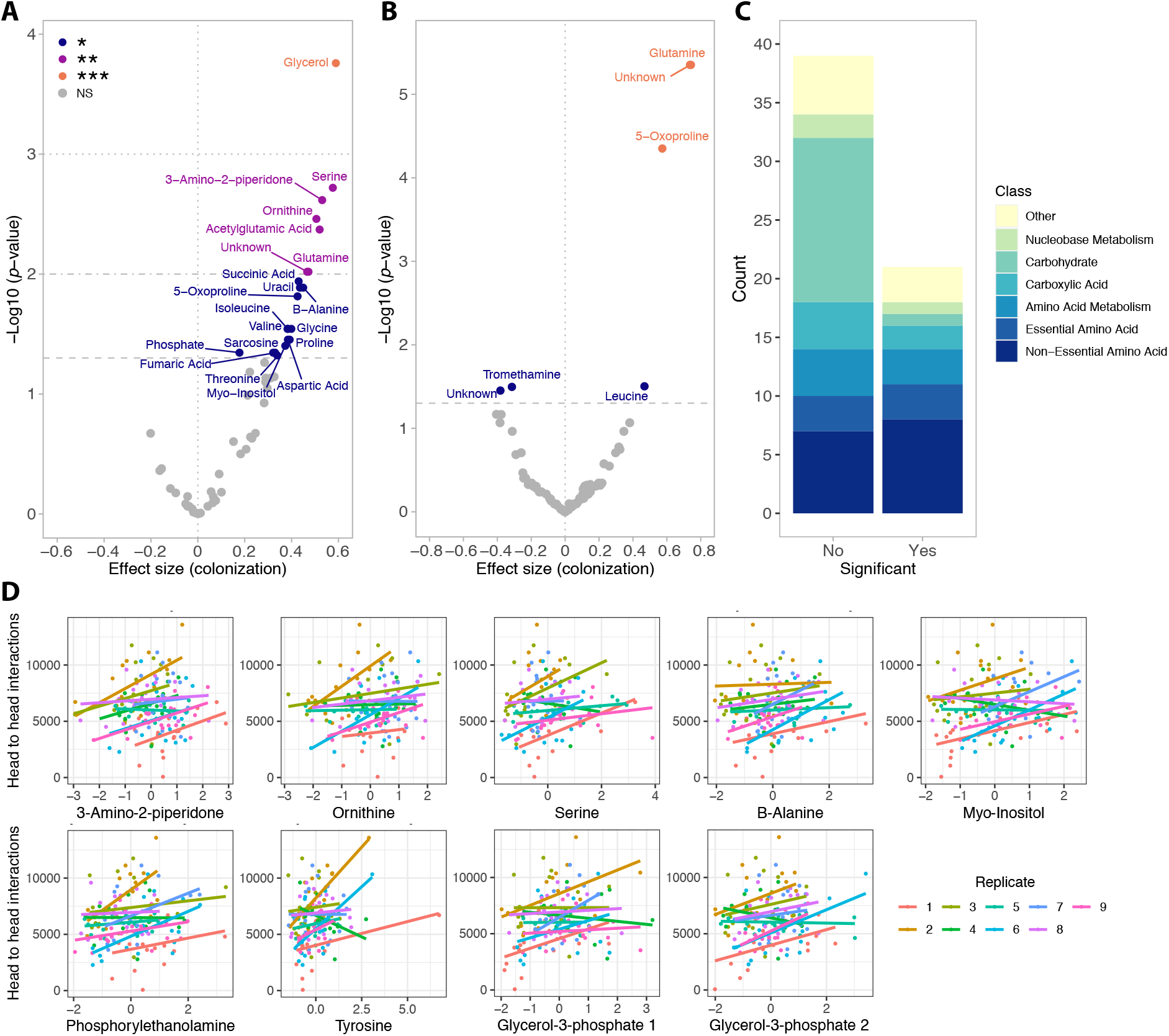
The gut microbiota increases the abundance of brain metabolites. Volcano plots present significance (-log10 (*P* value)) *versus* effect size of linear mixed effects models for all soluble metabolites identified in the brain (A) and hemolymph (B) of tracked bees. Positive effect sizes indicate metabolites that were more abundant in the brains of CL bees than in those of MD bees. *P* values were corrected for multiple testing with the BH method. *** *P<*0.001, ** *P<*0.01, * *P<*0.05, NS = not significant. (C) Stacked bars show the relative proportion of metabolites based on functional classification and plotted separately based on significance in differential abundance tests. (D) Regressions between metabolite abundance (z-score) and the number of head to head interactions of each bee for the nine metabolites that were significant predictors in linear mixed-effects models. Regression lines are colored by experimental replicate. The top row presents the metabolites that were also significantly differently abundant between MD and CL bees, the bottom row presents the four that were not.

Five of the 21 metabolites that were more abundant in brains of CL bees were significant predictors of the number of head to head interactions (Fig. 2D and Table S3; 3-amino-2-piperidone, ornithine, serine, B-alanine and myo-inositol; n=161, linear mixed-effects models fitted by REML with replicate as random effect, BH-adjusted *P<*0.05). Four other metabolites (phosphorylethanolamine, tyrosine and two identified as glycerol-3-phosphate) were also predictors of the rate of head to head interactions although their concentrations were not significantly different between CL and MD bees (Fig. 2D and Table S3). By contrast, none of the 76 hemolymph metabolites were significant predictors of the number of head to head interactions (Table S3; n=153, linear mixed-effects models fitted by REML with replicate as random effect, all BH-adjusted *P>*0.05). These findings suggest that the gut microbiota specifically increases the abundance of brain metabolites, which could be due to bacterial signals received from the gut or the direct transfer of microbial or dietary-derived metabolites from the gut to the brain. The latter seems likely for the three metabolites found to be more abundant in both the brain and the hemolymph of CL bees (a pattern consistent with transfer from the gut to the brain), and for essential amino acids, which the honeybee lacks the ability to produce (*33*).

We next investigated the effect of the gut microbiota on gene expression in the gut and three macro-regions of the brain that are broadly responsible for learning and memory (mushroom bodies, MB), perception of olfactory and gustatory stimuli (antennal lobe and subaesophageal ganglion, AL), and visual processing (optic lobes, OL) (Fig. 3A). As in the previous experiments, we reared sub-colonies (N=10) of either ∼20 CL or MD bees. From each sub-colony we randomly sampled a single bee for gene expression (to avoid cage effects, see Materials and Methods) and two-three additional bees for gut microbiota analyses. We included two additional treatments (also N=10) where bees were colonized with a synthetic community of 13 strains covering most phylotypes of the honeybee gut microbiota (CL_13; Table S4), or with only *Bifidobacterium asteroides*, which is thought to have neuromodulating potential (CL_Bifi; (*26*)). Seven days after inoculation, the gut microbiota clearly differed as expected between the four treatments (Figs. S5 and S6; PERMANOVA using Bray-Curtis dissimilarities calculated from a matrix of absolute ASV abundances: *F*_3,120_=50.80, *R*^2^=0.56, *P<*0.001). In the gut, a total of 4,988 bee genes (40% of the transcriptome) were differently expressed between MD and the three types of CL bees (See Materials and Methods; Figs. 3B, and S7 and Table S5). Colonization with only *B. asteroides* recapitulated a considerable subset of the changes associated with full colonization: more than a quarter (1267/4753) of the genes differentially expressed between MD and one or both of CL and CL_13 were also differently expressed between MD and CL_Bifi (Fig. S8). Moreover, only 15 genes were differentially expressed only between MD and CL_Bifi bees (Fig. S8).

**Figure 3:**
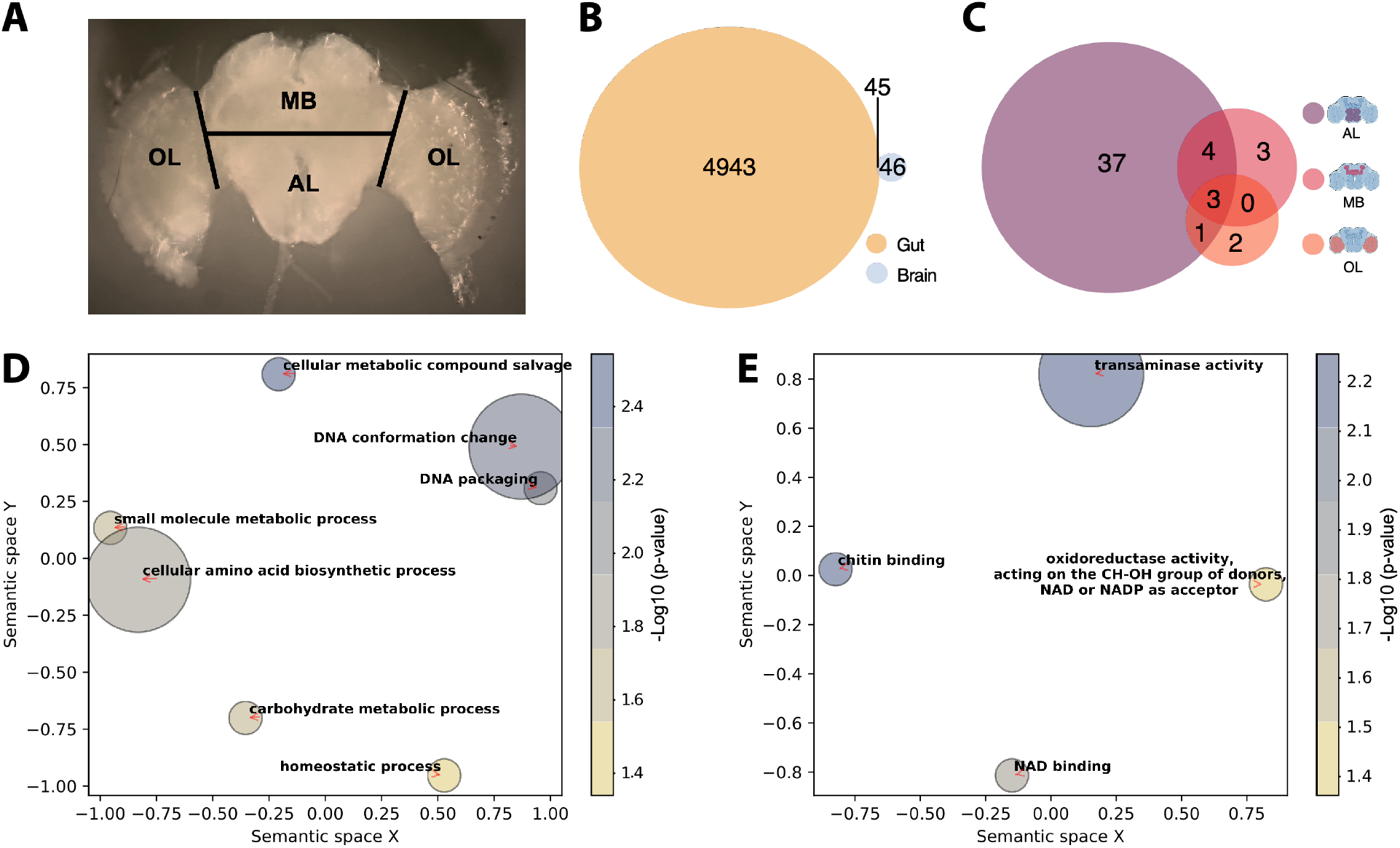
The gut microbiota alters gene expression in the gut and in the AL brain region. (A) Brain regions dissected for RNA sequencing. Black lines indicate the performed incisions. (B) Venn diagram reporting overlap in differentially expressed genes in the gut and brain across all contrasts of gut microbiota colonization treatments (CL, CL_13, and CL_Bifi) *versus* microbiota-depleted (MD) controls. (C) Venn diagram reporting overlap in brain-region-specific contrasts of all gut microbiota colonization treatments (CL, CL_13, and CL_Bifi) *versus* microbiota-depleted (MD) controls. Semantic similarity scatterplots summarize the list of enriched (D) Biological process and (E) Molecular function GO terms of all the 91 DEGs identified the brain. The scatterplots show GO terms as circles arranged such that those that are most similar in two-dimensional semantic space are placed nearest to each other. Circle color represents –log10 of enrichment *P* value.

In contrast to the widespread changes in the gut, the microbiota affected the expression of relatively few genes in the brain. Only 91 genes were differentially expressed between MD bees and bees of any of the three colonization treatments (Fig. 3B and Table S6). The proportion of these genes (45/91) that were also differentially expressed in the brain was greater than expected by chance (Fig. 3B; Hypergeometric test: representation factor = 1.23, *P*=0.047). The AL was the brain region with the greatest number of genes affected by the gut microbiota (Figs. 3C and S9 and Table S6). Consistent with our metabolomic analyses, the differentially expressed genes were mainly enriched for Gene Ontology terms related to amino acids and their metabolism (Fig. 3D and E and Table S7). Other enriched terms were related to the epigenetic regulation of chromosome packaging and conformation (Fig 3D and E and Table S7). The antennal lobes process information from the antennae, while the subaesophageal ganglion processes gustatory stimuli. Hence, our gene expression results together with our behavioral findings suggest that the gut microbiota increases social tendency by modulating chromatin accessibility and amino acid biosynthesis and metabolism in areas of the brain implicated in the perception of sensory stimuli. Previous work in mice also found that the gut microbiota affects amino acid metabolism in the host brain (*34*), suggesting that the mediation of the gut-brain axis via amino acid metabolism may be deeply conserved. In the brain, amino acids act as neurotransmitters, regulators of energy metabolism and neuromodulators, and imbalances are associated with neurodegeneration (*35*).

In conclusion, our study shows that the gut microbiota affects the rate of social interactions and the social network structure of honeybees. These behavioral differences are associated with important changes in gene expression and metabolite abundance in the brain. Our results demonstrate crosstalk between the gut microbiota and amino acid metabolism, particularly across the antennal lobes and the subaesophageal ganglion, the brain regions associated with perception of olfactory and gustatory stimuli (*36, 37*). Because changes in the rate and patterning of social interaction probably impact information and nutrient flow within colonies, our study highlights the importance of the gut microbiome for the complex social lives of honeybees.

## Materials and Methods

### Preparation of bacterial inocula to colonize microbiota-depleted bees

We produced three kinds of inocula: (i) a homogenate of five pooled guts of nurse bees collected from a single hive (CL treatment), (ii) an artificial community reconstituted from 13 cultured strains spanning the major phylotypes and SDPs (*26, 38*) of the honeybee gut microbiota (CL_13 treatment; Table S4), and (iii) an inoculum containing two cultured strains of *Bifidobacterium asteroides* (CL_Bifi treatment; Table S4).

To prepare the inoculum for the CL treatment, we collected five nurse bees from each of three hives and bead-beat their guts in 1 ml 1x PBS with 0.75 - 1 mm sterile glass beads using a FastPrep-24 5G homogenizer (MP Biomedicals) at 6 m/s for 45 s. We pooled the five gut homogenates by hive of origin, and plated a serial dilution of these pools from 10^−3^ – 10^−12^ onto BHIA, CBA + blood and MRSA + 0.1% L-cys + 2% fructose using the drop method (10 µl droplets). These plates were then incubated in both anaerobic and microaerophilic conditions to confirm bacterial growth prior to inoculations. To select the most pathogen-depleted of these three homogenates for subsequent colonizations, we performed diagnostic PCRs on lysates, using specific primers targeting known bee pathogens (*Nosema apis, Nosema ceranae*, trypanosomatids, *Serratia marcescens*, fungi, as well as *Bifidobacterium* as initial validation that homogenates contained live members of the core gut microbiota). Lysates were produced by mixing 50 µl of the homogenate with 50 µl lysis buffer, five µl proteinase K (20 mg/ml) and five µl lysozyme (20 mg/ml) and incubating these mixes for 10 min at 37 °C, 20 min at 55 °C and 10 min at 95 °C in a PCR machine. We then centrifuged these lysates for 5 min at 2000 g and used the supernatants as templates for PCR. We selected the homogenate that showed the least amplification of pathogen DNA.

For the CL_13 and CL_Bifi treatments, bacterial strains were inoculated from glycerol stocks and restreaked twice. Details on bacterial strains and culture conditions are reported in Table S4. We harvested bacterial cells and resuspended them in 1x PBS at an OD600 of 1. These suspensions were pooled in equal volumes in a falcon tube and pelleted by centrifugation at 4000 g for 5 min, after which we resuspended the pooled pellet in 1.5 ml PBS, added glycerol to the final concentration of 20% and stored the final CL_13 and CL_Bifi inocula at -80 °C.

### Automated behavioral tracking

Colonies of *Apis mellifera carnica* were reared at the University of Lausanne. Microbiota-depleted bees were produced as previously described (*26, 27*). Briefly, melanized dark-eyed pupae were individually extracted from capped brood cells with sterile forceps and placed in sterilized plastic containers lined with moist cotton. We performed nine experimental replicates of the automated behavioral tracking experiment. For each experimental replicate, we extracted four hundred pupae from one of nine different hives and placed them in 16 sterile plastic boxes in groups of 25. We then kept these pupae in an incubator at 70% RH and 34.5 °C in the dark. Three days later, we used superglue to affix 1.6 mm^2^ fiducial markers from the ARTag library (*39*) onto the thorax of all newly emerged workers that showed no sign of wing deformation. On the same day, we transferred these bees to each of 16 new cup-cages built with a sterile plastic cup placed on top of a 100 mm petri dish, and provided them with their treatment solutions. To do this, we added 100 µl droplets of either a gut homogenate (CL) diluted 1:1 in sugar water (SW) or a 1:1 PBS:SW solution as control (MD) to the petri dish. Pollen and sugar water were provided *ad libitum*. Two days later we pooled these bees according to their treatment group and transferred ca. 100 bees per treatment into two pairs of plexiglas boxes (22.5 cm length x 13.5 cm width) closed by transparent covers 1.5 cm above the floor and connected by a (50 cm length x 1.9 cm diameter) plastic tube (Fig. 1A and Supplementary Movie 1). These pairs of plexiglas boxes were hosted within separate climate-controlled chambers and monitored by a pair of tracking devices (all technical specifications and code available at: https://github.com/formicidae-tracker/). We defined a nest chamber by keeping one box under a constant 70% RH and 30 °C regime in the dark. In the foraging arena, climatic conditions cycled from 25°C and light during the day to 18°C and dark during the night (Fig. 1A). Transitions were initiated at 04:00 and 16:00, and programmed to last for four hours, during which the climate system performed a linear interpolation between the two states. Each box contained a trough filled with 1 g of pollen and three 2 ml vials filled with SW. These SW feeders were continuously replaced during the experiment. Bees were left to acclimatize in their boxes for a few hours, after which we conducted behavioral tracking from 00:00 to 08:00 on the same day of the subsequent week (a total of 152 hours). During this time, the x,y coordinates and orientation of each tag was recorded six times per second. At the end of each experiment, we counted and removed dead bees. We then scanned the tags to retrieve the identity of each bee, flash froze the bees in liquid nitrogen and stored them at -80 °C until further processing.

The tracking data were processed in FortStudio (https://github.com/formicidae-tracker/), where the body-length of each bee (front edge of clypeus – tip of abdomen; Fig. S10) was measured and polygons were drawn to define individual head and body regions (Fig. 1B). Data were subsequently processed using the R package FortMyrmidon (https://github.com/formicidae-tracker/). Contact events (i.e., the overlap of the body polygons) were saved along with the contact type (i.e., head to head, head to body, *etc*) and duration. Bees that interacted less than 2*SD below the mean interaction count of the sub-colony were excluded from all behavioral analyses. Average speed and standard deviation in speed were calculated from individual trajectories (time-calibrated x,y coordinates).

Statistical analyses were performed in R v4.1.0 (*40*). To assess the effect of the gut microbiota on behavioral variables (average values for each sub-colony) we ran paired *t* tests after checking that the differences between paired values were normally distributed using the Shapiro-Wilk normality test.

### Production of gnotobiotic bees for RNA-sequencing

We collected six boxes of 25 pupae from each of 10 hives and kept them in an incubator at 70% RH and 34.5 °C in the dark for two days. On the afternoon of the second day, we dissected one newly-emerged bee per box, homogenized their hindguts in 1 ml 1x PBS with 0.75 - 1 mm sterile glass beads using a Fast-Prep24 5G homogenizer (MP Biomedicals) at 6 m/s for 45 s. We then plated these homogenates on BHIA, CBA + blood and MRSA + 0.1% L-cys + 2% fructose growth media with the drop method and cultured them overnight in anaerobic and microaerophilic conditions. To minimize the risk of including contaminated bees in colonization experiments, the next day we excluded rearing boxes in which bacterial growth was observed for the tested bee, which led to the exclusion of two out of 60 boxes. Next, we transferred bees from each of the 58 remaining boxes into a corresponding sterile cage built with a 100 mm petri dish and an inverted sterile plastic cup of 3 dl.

Four cages belonging to each of the ten hives were randomly assigned to one of four treatments. Bees were either (i) kept microbiota-depleted (MD) or colonized with (ii) the gut homogenate (CL) inoculum (iii) the community of 13 strains (CL_13) inoculum, or (iv) the two strains of *Bifidobacterium asteroides* (CL_Bifi) inoculum. Colonizations were performed three days after pupae extraction. After thawing the inocula on ice, we diluted them in 1X PBS and 1:1 in sugar water (SW). We placed three droplets of 100 µl colonization suspensions at the bottom of the cages so that bees would be inoculated by physical contact with the suspension and trophallaxis with other bees. MD bees were given only a 1:1 PBS:SW solution (the extra cages that we produced for each hive were left MD to produce a surplus of these bees as a backup in case of contamination). Bees were provided with 1 g of sterile pollen and SW *ad libitum* and reared in an incubator at 70% RH and 30 °C in the dark.

One week post-treatment, we anesthetized bees on ice and dissected their guts excluding the honey stomach, which is generally colonized by environmental microbes that do not represent the core gut microbiota (*24, 41*). We then flash-froze the heads and guts and stored them in liquid nitrogen.

### Nucleic acid extraction from gut tissue

After having conducted the behavioral tracking experiment, we extracted DNA from the guts of 180 randomly selected bees (ten per replicate per treatment), for which we also performed metabolomics analyses of brain and hemolymph samples. We also performed one blank extraction (with no experimental tissue) per replicate (N=9) to identify and exclude laboratory reagents contaminants from 16S rRNA gene amplicon sequencing data (see below). The bees’ abdomens were thawed on ice, and guts were dissected and homogenized in a FastPrep-24 5G homogenizer (MP Biomedicals) at 6 m/s for 45 s in 360 µl ATL buffer and 40 µl proteinase K (20 mg/ml) containing ca. 100 µl of 0.1 mm Zirconia/Silica beads (Carl Roth). These homogenates were digested at 56 °C overnight, after which DNA was extracted from half of each homogenate using a Qiagen BioSprint 96 robot with the BioSprint DNA Blood Kit following the manufacturer’s instructions, including an RNase treatment step.

For the RNA-seq experiment, we randomly selected three-four bees per treatment per hive (N=121) for DNA extraction of gut samples. Guts were thawed on ice and homogenized in a FastPrep-24 5G homogenizer (MP Biomedicals) at 6 m/s for 45 s in 1 ml 1X PBS containing ca. 100 µl of 0.1 mm Zirconia/Silica beads (Carl Roth). Half of the volume of these homogenates was used for DNA extraction while the remaining homogenate from 40 of these bees (one randomly selected bee per treatment per hive from 40 independent cages, from which we also obtained brain RNA-seq data; see below) was used for RNA extraction. Nucleic acids were extracted with hot phenol protocols as previously described (*26*). We once more performed blank DNA extractions (with no experimental tissue) in parallel to control for laboratory reagent contaminations.

### Quantification of bacterial loads in the guts of gnotobiotic bees

We determined bacterial loads by qPCR using universal primers targeting the 16S rRNA gene as per Kešnerová *et al*. (*27*). qPCRs targeting the *Actin* gene (*27*) were used as controls of DNA quality. We also screened cDNA reverse-transcribed from gut RNA of the 40 bees that we selected for RNA-sequencing for the presence of Varroa destructor virus 1 (VDV-1) and deformed wing virus (DWV). There was no amplification of viral RNA from any of these samples. All qPCR reactions were carried out in 96-well plates on a StepOnePlus instrument (Applied Biosystems) following the protocols and using the primers reported in Kešnerová *et al*. (*26, 27*).

### 16S rRNA gene amplicon-sequencing

The V4 region of the 16S rRNA gene was amplified following the Illumina 16S metagenomic sequencing preparation guide (https://support.illumina.com/documents/documentation/chemistry_documentation/16s/16s-metagenomic-library-prep-guide-15044223-b.pdf) and the protocols and primers reported in Kešnerová *et al*. (*27*). Amplicon-sequencing was performed on an Illumina MiSeq sequencer at the Genomic Technology Facility of the University of Lausanne. Sequencing was done for 500 cycles, producing 2 × 250-bp reads.

### Analyses of 16 rRNA gene amplicon-sequencing data

We sequenced 16S rRNA gene amplicons from gut samples, bacterial inocula, negative PCR controls, and blank DNA extractions. We also included a mock community sample consisting of equal numbers of nine plasmids (pGEM®-T Easy vector; Promega) containing eight 16S rRNA gene sequences from honeybee gut symbionts and one from *E. coli*, which we used as internal standard to verify consistency between MiSeq runs. Raw sequencing data (deposited at the SRA Database under Accession no. PRJNA792398) were quality-controlled with FastQC (http://www.bioinformatics.babraham.ac.uk/projects/fastqc/) and primer sequences were removed with Cutadapt (*42*). We then continued the analysis using the Divisive Amplicon Denoising Algorithm 2 (DADA2) package v.1.20.0 (*43*) in R. All functions were run using the recommended parameters (https://benjjneb.github.io/dada2/tutorial.html) except that at the filtering step we truncated the F and R reads after 232 and 231 bp, respectively. We then set randomize=TRUE and nbases=3e8 at the learnErrors step. We used the SILVA database (version 138) to classify the identified amplicon-sequence variants (ASVs). To complement the taxonomic classification based on the SILVA database, sequence variants were further assigned to major phylotypes of the bee gut microbiota as previously defined (*27*). Any unclassified ASV was removed with the “phyloseq” package version 1.36.0 (*44*), using the “subset taxa” function. We then used both the “prevalence” and “frequency” methods (method = “either”) in the R package “decontam” v.1.12.0 (*45*) to identify and remove contaminants introduced during laboratory procedures, using the negative PCR controls and the blank samples as reference.

### Analyses of combined 16S rRNA gene amplicon-sequence and qPCR data

To calculate absolute bacterial abundances of each ASV, the proportion of each ASV in each sample was multiplied by the total 16S rRNA gene copy number of each sample as measured by qPCR (*27*). To assess differences in community structure between treatments we ran ADONIS tests after calculating Bray-Curtis dissimilarities with the absolute ASV abundance matrix.

### Extraction of metabolites from tracked bees

We analyzed soluble metabolites in the brain and hemolymph from the random subset of 180 tracked bees for which we also analyzed the gut microbiota. CL bees in this subset engaged in a greater number of head to head interactions than MD bees, consistent with our global analysis (Fig. S11; linear mixed-effects model fitted by REML with experimental replicate as random effect: n=174, *F*_1,164_=12.15, *P<*0.001). Brains were dissected from frozen bees, weighed on a microbalance, and refrozen at -80 °C until extraction. Hemolymph (1 µl) was taken from the thorax of thawed bees and refrozen at -80 °C until extraction. Individual brain and hemolymph samples were extracted following a modified Bligh and Dyer protocol (*46-48*). Frozen brain tissue was ground with a motorized pestle for 30 s in 100 µl of chilled (4:1) analytical grade methanol:ddH_2_O with 1 mM norluecine (Sigma Aldrich) standard. Hemolymph was extracted in the same mixture, omitting the tissue-grinding step. Samples were then extracted in a thermomixer (10 min, 2000 rpm, 4 °C) and centrifuged (5 min, 15000 rcf, 4 °C). Supernatant was transferred to a new tube and kept chilled at -20 °C, while 250 µl of cold (1:1) chloroform:methanol (Sigma Aldrich) was added to the sample. Samples were again extracted in the same manner, and the supernatants combined. Phase separation was achieved with 200 µl ddH_2_O, followed by a fast vortex and centrifuge step. The top aqueous layer was removed and dried in a speedvac concentrator overnight at ambient temperature. The sample was derivatized with 50 µl of 20 mg/ml methoxyamine hydrochloride in pyridine (Sigma Aldrich), for 90 min at 33 °C followed by silylation with 50 µl of MSTFA (Sigma Aldrich) for 120 min at 45 °C.

### GC-MS analysis of metabolites

Samples were analyzed on an Agilent 8890-5977B GC-MSD equipped with a Pal3 autosampler that injected 1 µl of sample onto a VF-5MS (30 m x 0.25 mm x 0.25 mm) column. The samples were injected with a split ratio of 15:1, helium flow rate of 1 ml/min and inlet temperature of 280 °C. The temperature was held for 2 min at 125 °C, raised at 3 °C/min to 150 °C, 5 °C/min to 225 °C, and 15 °C/min to 300 °C and held for 1.3 min. The MSD was run in scan mode from 50-500 Da at a frequency of 3.2 scan/s. Spectral deconvolution and compound identification was performed with Masshunter Workstation Unknown Analysis software (Agilent) and the NIST 2017 MS library. Best hits of compound identity are reported for spectra with a match factor greater than 85%. Identified metabolites were then manually mapped to metabolic pathways in the KEGG PATHWAY Database. Analyte abundances were calculated using the MassHunter Workstation Quantitative Analysis software (Agilent).

### Metabolomics analysis

Raw metabolite abundances were normalized to the internal standard and then to the sample mass (brains only). Low-quality samples and samples with an ISTD response *<* or *>* two SD from the batch mean were removed from the datasets. The normalized abundances were then transformed to z-scores. The impact of colonization on metabolite abundance was then calculated using a mixed integer linear model using the *lmm2met* package in R (*49*). Colonization was treated as a fixed effect, while the nine different experimental batches were treated as a random effect. One global batch term was used, as each step in the extraction and analysis pipeline was performed in the same paired batch fashion as in the automated behavioral tracking experiment. The significances of the effect sizes were calculated using a likelihood-ratio test and adjusted using the Benjamini–Hochberg (BH) procedure. We next performed separate linear mixed-effects models between the abundance (z-score) of each metabolite (independent variable) and the number of head to head interactions of each bee (dependent variable). We considered the different experimental batches as a random effect and adjusted for multiple testing with the BH method.

### RNA-sequencing of gut and brain tissues

For RNA-sequencing, we randomly selected one bee per treatment per hive (40 total bees), so that all samples were independently reared in separate cages (no cage or hive effect). We sequenced RNA from the gut and brain of each individual. The heads were moved from liquid nitrogen into RNAlaterICE (Life Technologies) in a petri dish placed onto a metal plate chilled on ice. We immediately dissected the brain with sterile forceps, after carefully removing the hypopharyngeal glands, compound eyes and ocelli and further dissected the brain into three macro-regions by performing a horizontal incision across the midbrain through the posterior protocerebral lobe and two oblique incisions to separate the optic lobes from the rest of the brain (Fig. 3A), using needles. The resulting regions were: the optic lobes (OL), the mushroom body region (MB), and the lower part of the midbrain, containing the antennal lobes and the subesophageal ganglion (AL). RNA extractions of brain regions were performed with the Arcturus PicoPure RNA Isolation Kit (Applied Biosystems) according to the manufacturer’s specifications, including a DNase treatment (Qiagen) to remove genomic DNA. Brain-region samples were transferred to the kit’s incubation buffer and homogenized for 30 s with a motorized pestle.

The quality of both brain and gut RNA extractions was verified using a Fragment Analyzer (Advanced Analytical). RNA-sequencing libraries were prepared with the KAPA stranded mRNA kit (Roche) following the manufacturer’s protocol, except that we appended TruSeq unique dual indexes (UDIs, Illumina) instead of the adapters provided by the kit to better control for index hopping during sequencing. We always performed RNA extractions and library preparations for all bees from each hive/experimental replicate at the same time so as to only have one combined batch factor to control for. However, four bees had to be reprocessed as one of the tissues failed at library preparation. Hence, we accounted for an 11th RNA extraction / library preparation batch during analysis. Each sample was sequenced twice in separate sequencing lanes on a HiSeq 4000 sequencer (Illumina) at the Genomic Technology Facility of the University of Lausanne, producing single-end 150 bp reads.

### RNA-sequencing data analyses

Read quality was assessed with FastQC (http://www.bioinformatics.babraham.ac.uk/projects/fastqc/). We used Trimmomatic (*50*) to remove adapters and low-quality bases with the following parameters: LEADING: 10 (trim the leading nucleotides until quality *>* 10), TRAILING: 10 (trim the trailing nucleotides until quality *>* 10), SLIDINGWINDOW: 4:20 (trim the window of size four for reads with local quality below a score of 20), and MINLEN: 80 (discard reads shorter than 80 bases). Reads were then aligned with STAR v.2.5.4b (*51*) to the honeybee genome (*Apis mellifera* assembly HAv3.1 (*52*)). The two bam files belonging to each sample were merged with Samtools merge (*53*). Mapped reads were then converted into raw read counts with the htseq-count script (http://www.huber.embl.de/users/anders/HTSeq/doc/count.html). Two gut samples and four brain region samples (two OL, one AL, one MB) were not included in down-stream analyses because they either failed during library preparation or represented clear outliers, with less than 10% of reads mapping to the honeybee genome. We used the filterByExpr function in edgeR (*54*) to filter out genes that were not represented by at least 20 reads in a single sample. We then used the *Limma* Bioconductor package (*55*) for analyses of differential expression. For the gut we used the formula 0 + Treatment + Batch, whereas for the brain we used the formula 0 + group + Batch, where “group” represented every possible combination of brain region and treatment group and “Batch” represented the different experimental and RNA-seq library preparation batches. We accounted for the random effect of sampling multiple brain regions from the same individuals using the *duplicateCorrelation* function. The three different brain regions showed very distinct patterns of gene expression, indicating the precise dissection of the brain and quantification of region-specific gene expression (Fig. S12). We therefore performed the desired contrasts between brain regions and treatments, overall and within each brain region independently. *P* values of differential expression analyses were corrected for multiple testing with a false discovery rate (FDR) of 5%.

To perform Gene Ontology (GO) enrichment analyses we retrieved GO terms using biomaRt (amellifera_eg_gene dataset; (*56*)). We used a hypergeometric test implemented in the R Bioconductor package *GOstats* v.2.58.0 (*57*) to evaluate the differentially expressed gene lists for GO term associations, using the full genome as background and retaining GO terms with *P<*0.05. *GOFigure!* (*58*) was subsequently used to reduce redundancy in significant GO terms and summarize results by semantic similarity, using a similarity threshold of 0.8.

## Acknowledgements

We would like to thank Christine La Mendola and Catherine Berney for their technical support with RNA extraction and library preparation of honeybee brain samples, Théodora Steiner for continuous support in the laboratory, Matthias Rüegg and Alexandre Tuleu for technical assistance with the automated tracking system and Jule Wermerssen for the bee drawing in Fig. 1A. The two equally contributing senior authors flipped a coin to determine last authorship. This work was funded by the University of Lausanne, the European Union’s Horizon 2020 research and innovation programme under the Marie Skłodowska-Curie grant agreement BRAIN (no. 797113) to Joanito Liberti, by a HFSP Young Investigator grant (RGY0077/2016), an ERC Starting Grant (MicroBeeOme), and a Swiss National Science Foundation grant (31003A 160345) to Philipp Engel, and by a ERC Advanced Grant (resiliANT, no. 741491) to Laurent Keller.

## Author contributions

JL, PE and LKel conceived and designed the study. JL, PE and LKel acquired fundings. PE and LKel supervised the research. JL and TK performed the automated behavioral tracking experiment. TK performed automated behavioral tracking data analyses with assistance from JL and TOR. JL performed statistical analyses. JL performed microbiological preparations and gnotobiotic manipulations with assistance from LK, TK, AC and ETF. JL extracted DNA and JL and LK performed qPCR analyses. JL performed amplicon-sequencing and data analyses. JL performed gut and brain dissections and hemolymph collection. AQ performed metabolite extractions, GC-MS runs and metabolomics data analyses with assistance from JL. JL extracted RNA and analyzed RNA-sequencing data. LK performed RNA-sequencing library preparations. JL, TK and AQ plotted the graphs. JL, TK, PE and LKel drafted the manuscript. All authors contributed to interpreting the data and editing subsequent drafts of the manuscript.

## Competing interests

Authors declare no competing interests.

## Data and materials availability

Raw RNA-sequencing data have been deposited in NCBI’s Gene Expression Omnibus and are accessible through GEO Series accession number GSE192784 (https://www.ncbi.nlm.nih.gov/geo/query/acc.cgi?acc=GSE192784), while raw amplicon-sequence data are available on Sequence Read Archive (SRA) under accession PRJNA792398. Raw data tables, metadata and codes are available on GitHub at https://github.com/JoanitoLiberti/The-gut-microbiota-affects-the-social-network-of-honeybees. Additional input files required to reproduce the automated behavioral tracking analyses are available on Zenodo at: https://doi.org/10.5281/zenodo.5797980.

## Notes

https://github.com/JoanitoLiberti/The-gut-microbiota-affects-the-social-network-of-honeybees

https://doi.org/10.5281/zenodo.5797980

### Competing Interest Statement

The authors have declared no competing interest.

