## Supplementary figures and captions for "The gut microbiota affects the social network of honeybees"

.

**This PDF file includes:**

Figs. S1 to S12
Captions for Tables S1 to S7
Caption for Movie S1

**Other Supplementary Materials for this manuscript include the following:**

Tables S1 to S7
Movie S1

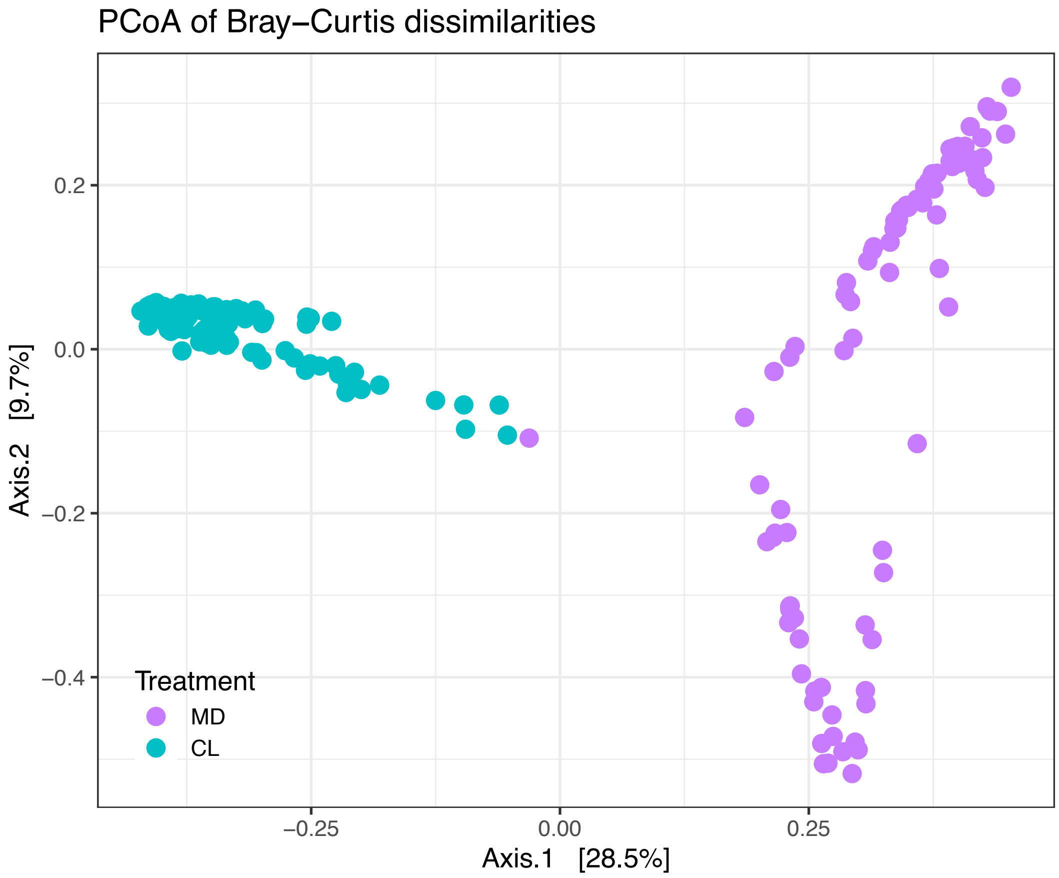


**Fig. S1.**

Principal Coordinate Analysis of Bray-Curtis dissimilarities between gut microbiota profiles in the automated tracking experiment. The ordination was performed on Bray-Curtis dissimilarities calculated from a matrix of absolute bacterial abundances of each amplicon-sequence variant (ASV) in each sample. This was obtained by multiplying the relative proportion of each ASV in each sample by the total number of 16S rRNA gene copies in the sample.

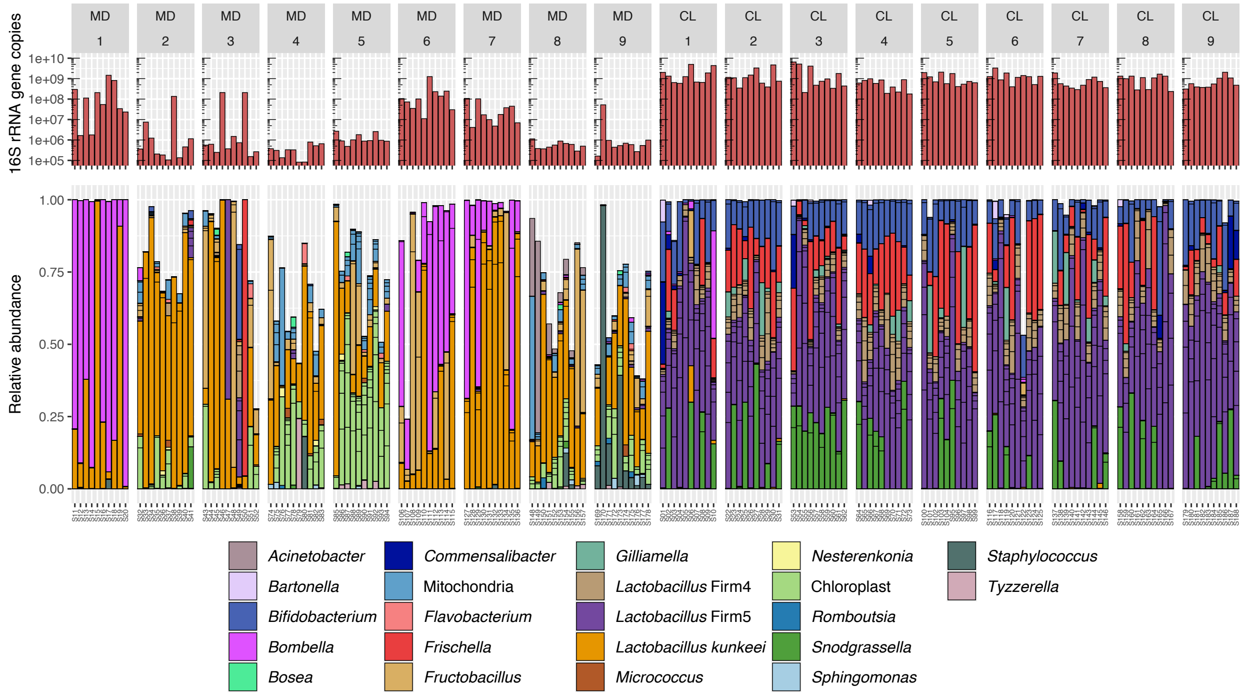


**Fig. S2.**

Bacterial loads and microbiota composition in the guts of bees in the automated tracking experiment. The upper barplots depict the number of 16S rRNA gene copies measured by qPCR with universal bacterial primers. Lower stacked bars indicate the relative abundance of community members. Multiple ASVs can have the same classification (color) and are separated by horizontal ticks. For ease of visualization, the stacked bars show only ASVs that had a minimum of 1% relative abundance in five samples.

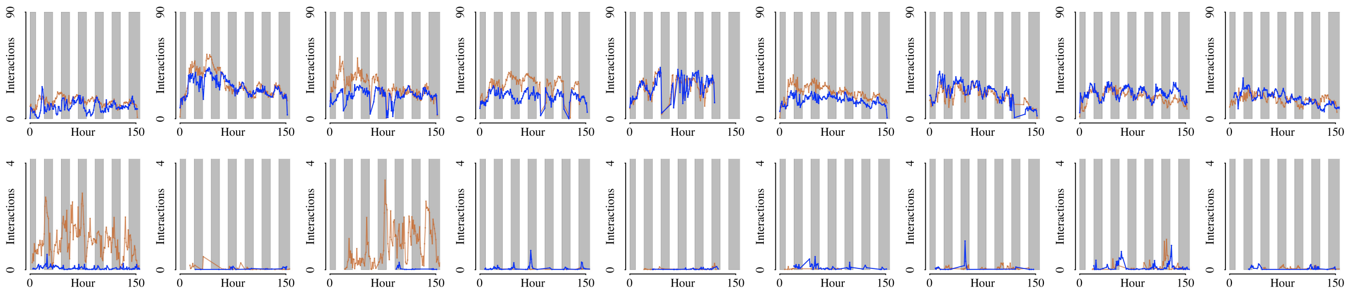


**Fig. S3.**

Line plots showing the number of head to head interactions per bee per hour in the automated behavioral tracking experiment. Columns correspond to experimental replicates. Top row = nest arena; bottom row = foraging arena. Brown lines = CL sub-colonies; blue lines = MD sub-colonies. Background bars show night (gray) and day (white). The expected circadian pattern of interaction frequency is apparent.

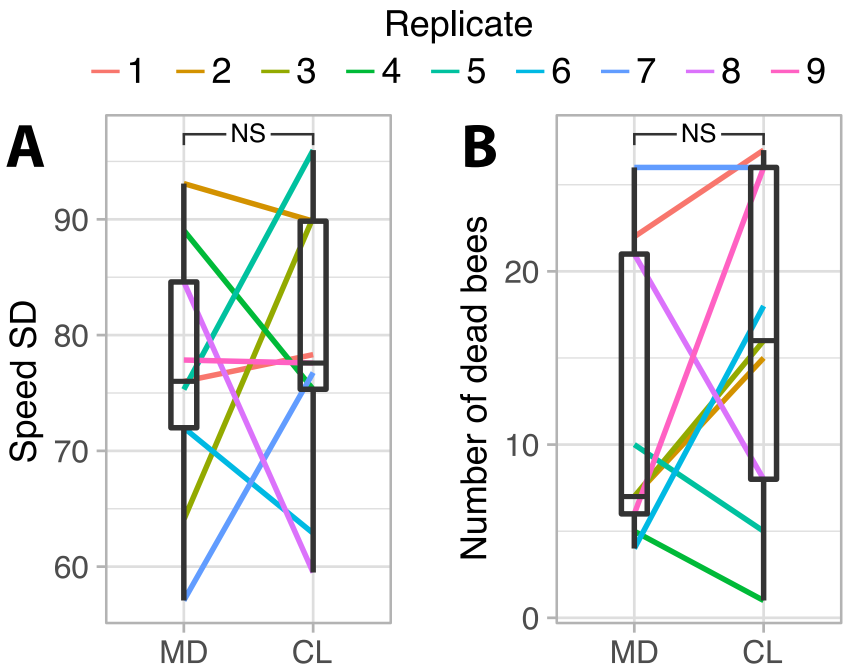


**Fig. S4.**

Standard deviation of speed (A) and mortality of tracked bees (B). Lines connect paired colonies in each experimental replicate. Boxplots show the median and first and third quartiles, while upper and lower whiskers report largest and lowest values within 1.5 times the interquartile ranges above and below the 75th and 25th percentiles, respectively. NS = not significant.

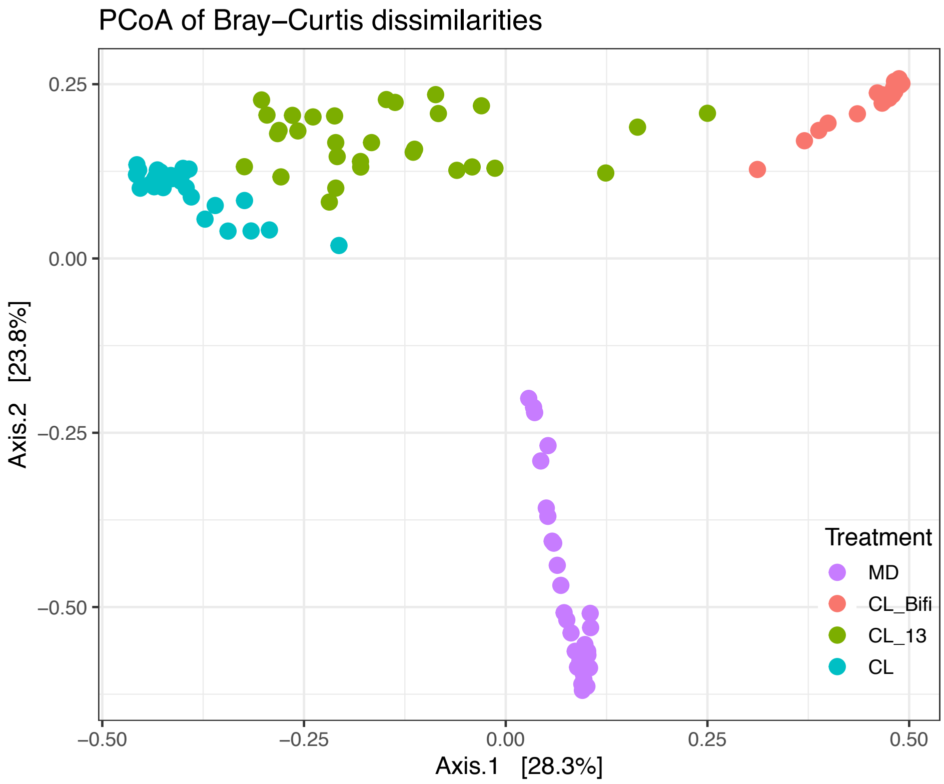


**Fig. S5.**

Principal Coordinate Analysis of Bray-Curtis dissimilarities between gut microbiota profiles in the RNA-sequencing experiment. Bray-Curtis dissimilarities were calculated from the absolute bacterial abundances of each ASV in each sample.

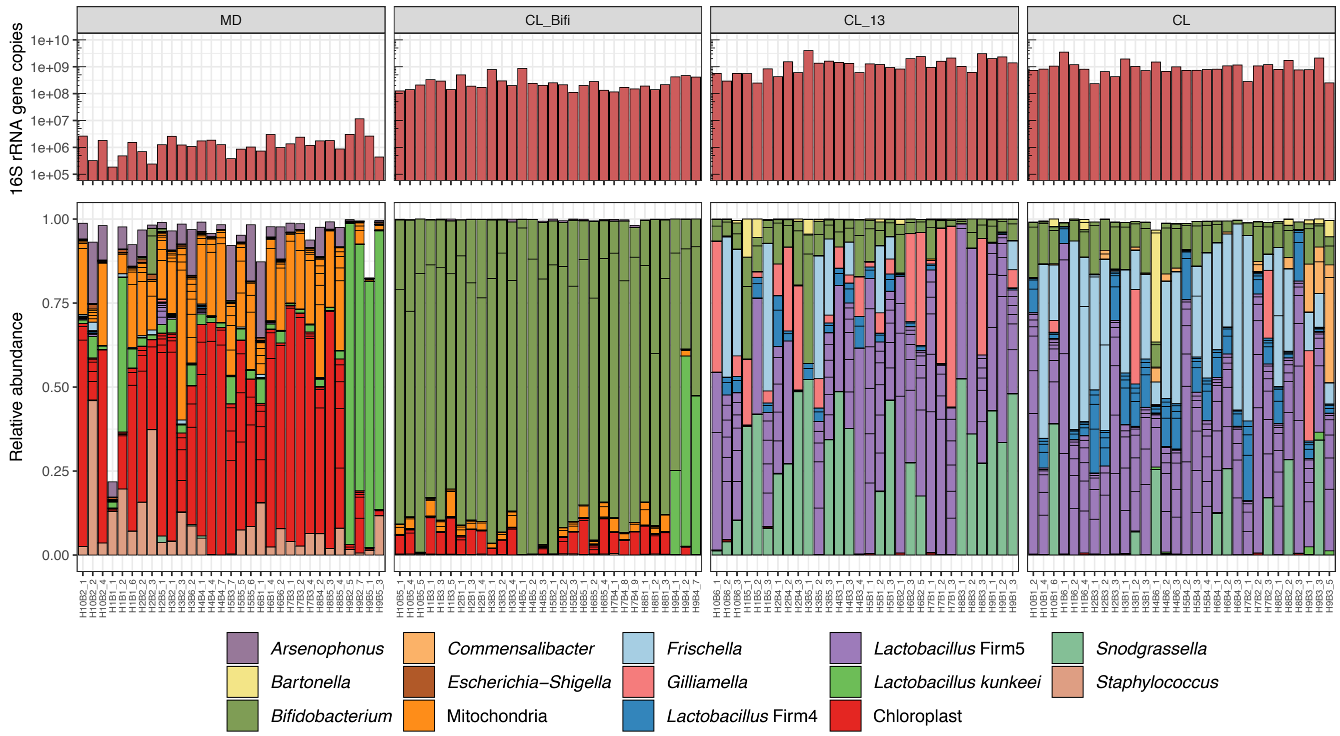


**Fig. S6.**

Bacterial loads and microbiota composition in the guts of bees from the RNA-sequencing experiment. The upper barplots depict the number of 16S rRNA gene copies measured by qPCR with universal bacterial primers. Lower stacked bars indicate the relative abundance of community members. Note that multiple amplicon-sequence variants (ASVs) can have the same classification (color) and are separated by horizontal ticks. For ease of visualization, the stacked bars show only ASVs that had a minimum of 1% relative abundance in two samples.

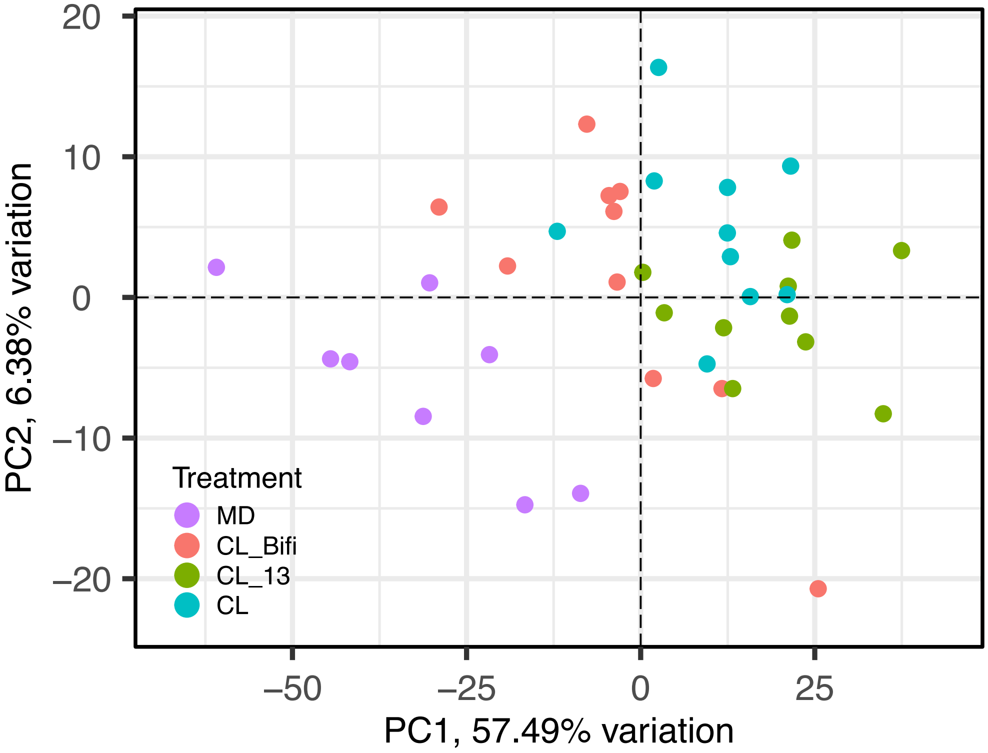


**Fig. S7.**

Principal Component Analysis of differentially expressed genes in honeybee gut samples. The ordination clusters the samples based on the expression (trimmed mean of M values (TMM) normalized counts) of 4,988 DEGs identified in contrasts of colonized treatments and microbiota-depleted controls. Samples are color-coded by gut microbiota treatment group.

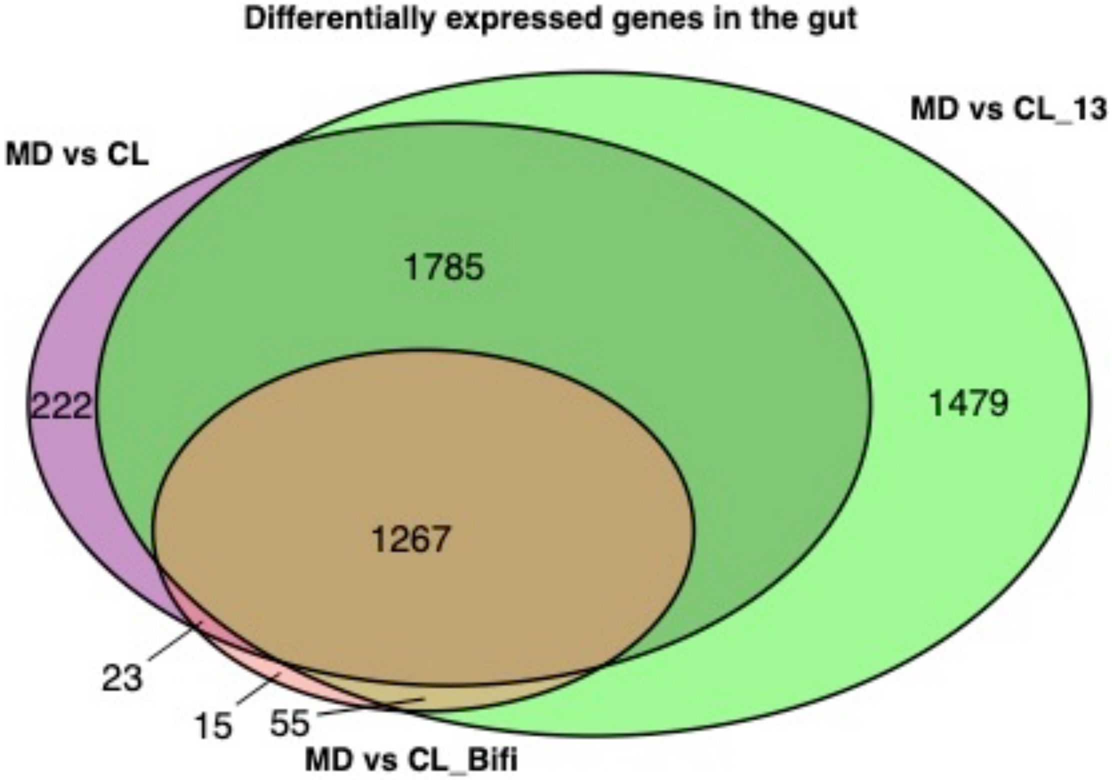


**Fig. S8.**

Venn diagram of DEGs in the gut. The diagrams report overlap in differentially expressed genes between contrasts of colonized treatments and microbiota-depleted controls in the gut. Note that additional comparisons between MD vs. both CL_13 and CL and between MD and all colonization treatments combined (CL_13, CL and CL_Bifi) have been omitted here for ease of visualization. See Table S5 for complete DEG lists.

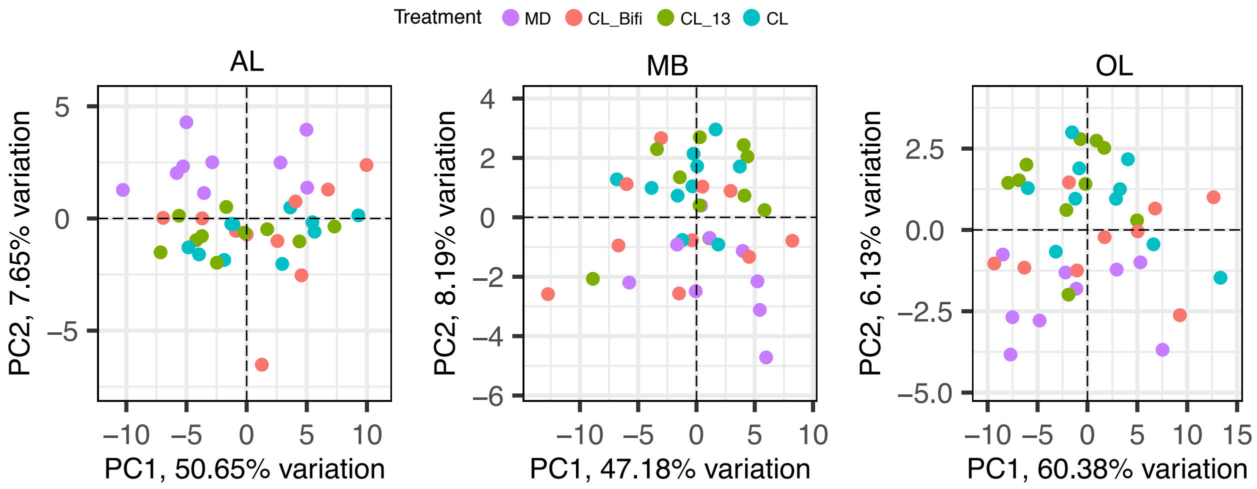


**Fig. S9.**

Principal Component Analyses of brain-region-specific expression of genes altered by the honeybee gut microbiota. The ordinations cluster samples based on the expression (TMM-normalized counts) of the 91 differentially expressed genes identified across whole-brain and region-specific contrasts of all colonized treatments against microbiota-depleted controls. Samples are color-coded by gut microbiota treatment group. AL = antennal lobes and subaesophageal ganglion, MB = mushroom bodies, OL = optic lobes.

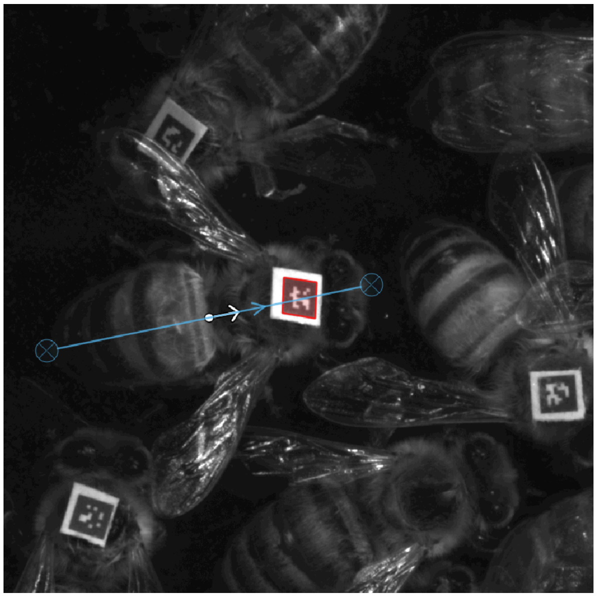

**Fig. S10.**

Example of the post-processing procedure to determine the orientation of a tracked bee. In FortStudio, a line was drawn from the tip of the abdomen to the front edge of the clypeus to derive the orientation of the tag relative to the body of the bee.

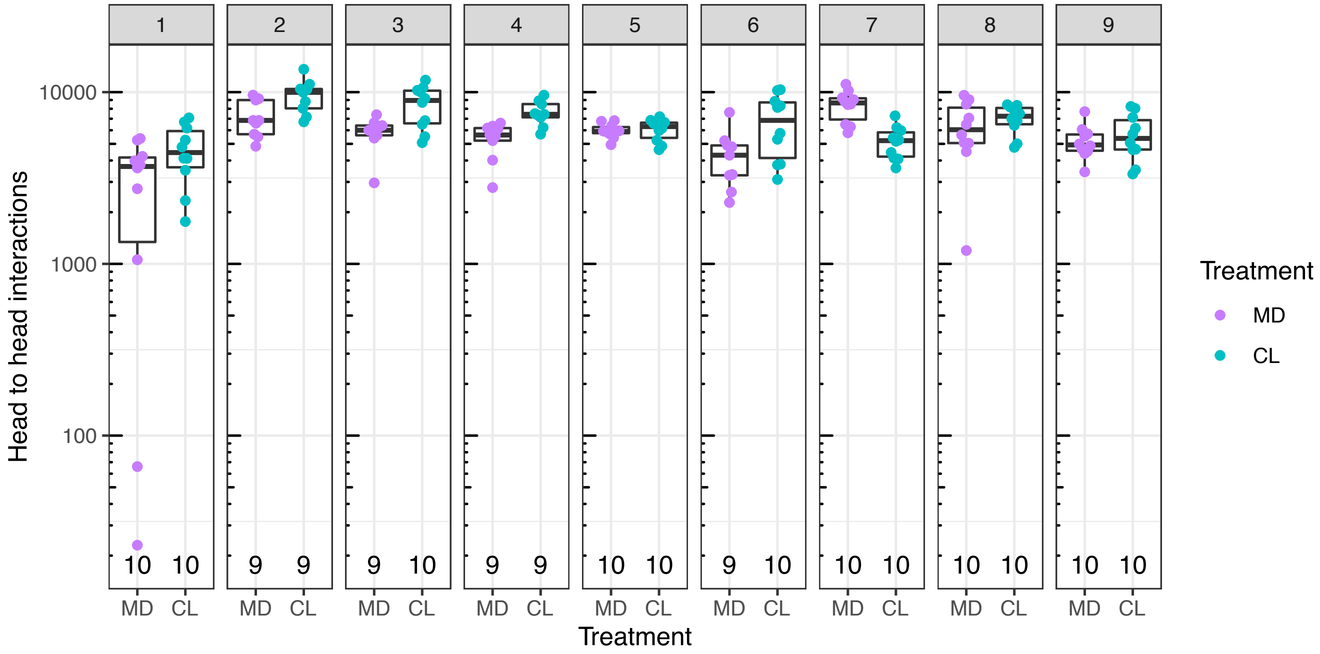


**Fig. S11.**

Social interactions in a subset of tracked bees by gut microbiota treatment group and experimental replicate. The plot shows the number of head to head interactions of the tracked bees for which we also obtained gut microbiota and metabolome data. For six of these 180 bees the number of head to head interactions could not be retrieved due to deterioration of the tags at the end of the week of tracking.

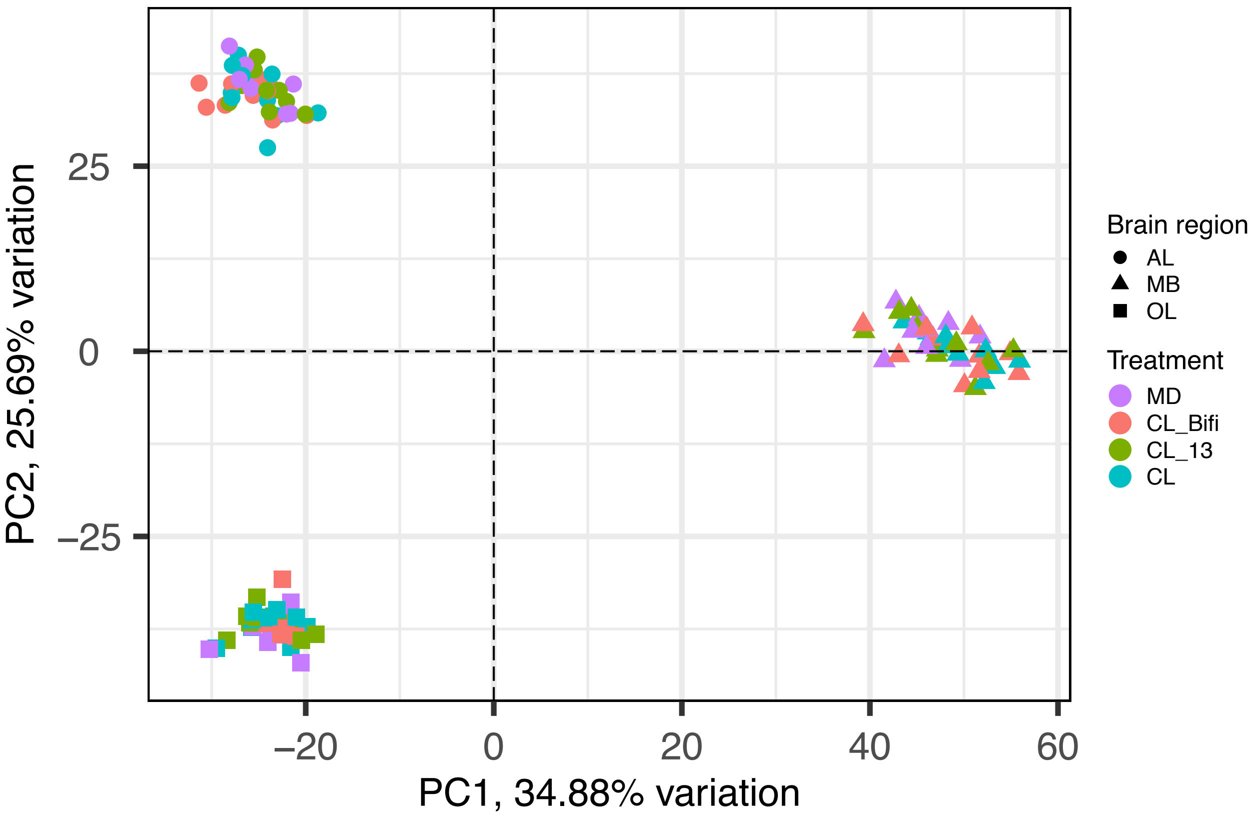


**Fig. S12.**

Principal Component Analysis of overall gene expression of brain samples. The ordination clusters samples based on the expression (TMM-normalized counts) of 10,493 genes retained after filtering out those with low expression and removing the experimental batch effect. Color indicates gut microbiota treatment group and shape indicates the different brain regions. AL = antennal lobes and subaesophageal ganglion, MB = mushroom bodies, OL = optic lobes.

**Table S1**. **(separate file)**
Results of differential metabolite abundance between gut microbiota treatments in the automated behavioral tracking experiment. Brain and hemolymph results are presented in separate sheets. For each metabolite the table shows the effect size and *P* value of the colonization treatment and the injection order in the GC-MS, as well as the BH-adjusted *P* values for the colonization treatment.

**Table S2**. **(separate file)**
Metabolite retention time, library identification, and functional categorization of the metabolites detected in the brain and hemolymph of bees in the automated behavioral tracking experiment, showed in separate sheets.

**Table S3**. **(separate file)**
Results of linear mixed effects models between abundance (z-scores) of individual metabolites and the number of head to head interactions of individual bees in the automated behavioral tracking experiment. For each metabolite, separate columns report the *F* value, the *P* value and the adjusted *P* value after BH-correction. Results for brain and hemolymph are shown in separate sheets.

**Table S4**. **(separate file)**
Identity and culture conditions of the strains used to produce the CL_13 and CL_Bifi inocula.

**Table S5**. **(separate file)**
Results of differential gene expression analysis of gut samples. Separate sheets report the lists of differentially expressed genes of different pair-wise comparisons.

**Tables S6**. **(separate file)**
Results of differential gene expression analysis of brain samples. Separate sheets report the lists of differentially expressed genes of different pair-wise comparisons.

**Table S7**. **(separate file)**
Results of gene ontology enrichment analysis of the list of 91 DEGs identified in the brain.

**Supplementary Movie 1**. **(separate file)**
Monitoring of social interactions under an automated behavioral tracking system. The video shows the nest box of one sub-colony. In this video orange lines connect bees whenever any kind of interaction occurs: body to body, head to head or head to body. Playback speed is 4x the actual speed.
